# Discovering human transcription factor interactions with genetic variants, novel DNA motifs, and repetitive elements using enhanced yeast one-hybrid assays

**DOI:** 10.1101/459305

**Authors:** Shaleen Shrestha, Jared Allan Sewell, Clarissa Stephanie Santoso, Elena Forchielli, Sebastian Carrasco Pro, Melissa Martinez, Juan Ignacio Fuxman Bass

**Author notes:** Correspondence: Dr. Juan Fuxman Bass. these authors contributed equally to this work (Shaleen Shrestha, Jared Allan Sewell).

## Abstract

Identifying transcription factor (TF) binding to noncoding variants, uncharacterized DNA motifs, and repetitive genomic elements has been technically and computationally challenging. Current experimental methods, such as chromatin immunoprecipitation, generally test one TF at a time, and computational motif algorithms often lead to false positive and negative predictions. To address these limitations, we developed two approaches based on enhanced yeast one-hybrid assays. The first approach interrogates the binding of >1,000 human TFs to repetitive DNA elements, while the second evaluates TF binding to single nucleotide variants, short insertions and deletions (indels), and novel DNA motifs. Using the first approach, we detected the binding of 75 TFs, including several nuclear hormone receptors and ETS factors, to the highly repetitive Alu elements. Using the second approach, we identified cancer-associated changes in TF binding, including gain of interactions involving ETS TFs and loss of interactions involving KLF TFs to different mutations in the *TERT* promoter, and gain of a MYB interaction with an 18 bp indel in the *TAL1* super-enhancer. Additionally, we identified the TFs that bind to three uncharacterized DNA motifs identified in DNase footprinting assays. We anticipate that these approaches will expand our capabilities to study genetic variation and under-characterized genomic regions.

## INTRODUCTION

The study of transcription factor (TF) binding to different genomic regions, and how TF binding is affected by noncoding variants, is critical for understanding the mechanisms by which gene expression is controlled in normal and pathogenic conditions (Gerstein et al. 2012; Fuxman Bass et al. 2015). Chromatin immunoprecipitation followed by next-generation sequencing (ChIP-seq) has been instrumental in identifying the genomic regions occupied by a TF and for studying TF function (Robertson et al. 2007; Gerstein et al. 2012). However, it has been challenging to use ChIP-seq to address many central human functional genomics problems such as determining whether disease-associated single nucleotide variants (SNVs) and short insertions/deletions (indels) in noncoding regions alter TF binding, as well as identifying TFs that bind to specific DNA motifs and repetitive genomic DNA elements.

Experimentally determining whether TF binding is altered by genomic variants associated with genetic diseases or cancer has been challenging as this cannot be performed using ChIP-seq without *a priori* TF candidates. This is because all ∼1,500 human TFs would have to be evaluated individually, and because samples from the appropriate tissues and conditions from healthy and sick individuals need to be obtained and compared (Fuxman Bass et al. 2015; Gan et al. 2018). Thus, the most widely used approach to prioritize TFs consists of using known DNA binding specificities (available for ∼50% of human TFs (Weirauch et al. 2014)) and motif search algorithms such as FIMO, BEEML-PWM or TFM-pvalue to compare predicted TF binding between the different noncoding alleles (Touzet and Varre 2007; Grant et al. 2011; Zhao and Stormo 2011; Weirauch et al. 2014; Rheinbay et al. 2017). However, this approach often results in multiple false positive and false negative predictions given that: 1) DNA motifs are missing for nearly half of the known human TFs (Weirauch et al. 2014), 2) most predicted DNA motifs in the genome are not occupied by the TF *in vivo* (Zia and Moses 2012), 3) multiple genomic regions occupied by TFs do not contain the corresponding TF binding sites (Gheorghe et al. 2018), and 4) sequence preferences determined using naked DNA may be different than for nucleosomal DNA (Talebzadeh and Zare-Mirakabad 2014; Zhu et al. 2018).

Many genomic regions identified by DNase I or ATAC-seq footprinting studies are occupied by unidentified proteins, which in many cases are bound in a sequence-specific manner (Neph et al. 2012; Ramirez et al. 2017). Indeed, using genome-wide DNase I footprinting, the ENCODE Project identified 683 de novo motifs, 289 of which could not be matched to any TF based on known DNA-binding specificities (Neph et al. 2012). Due to the lack of TF candidates, it is nearly impossible to use ChIP to identify the TFs interacting with these genomic sites, as hundreds of TFs would need to be tested in each of the cell lines where the footprints were found. Thus, most of the novel DNA motifs derived from DNase I footprinting remain uncharacterized.

More than half of the human genome is comprised of tandem or interspaced repetitive DNA elements, many located within promoter, enhancer, and silencer sequences (Lander et al. 2001). TF binding to these repetitive genomic elements is challenging to study by ChIP-seq, not only because hundreds of TFs need to be assayed, but also because repetitive DNA sequences are difficult to map to the reference genome and are thus often filtered out in most bioinformatics analysis pipelines (Rozowsky et al. 2009; Chung et al. 2011). This is particularly true for highly repetitive DNA elements such as the Alu short interspaced nuclear elements, which are present in more than one million copies in the human genome (Deininger 2011).

Enhanced yeast one-hybrid (eY1H) assays provide an alternative approach to ChIP-seq, where physical interactions between TFs and DNA regions are tested in the milieu of the yeast nucleus using reporter genes (Reece-Hoyes et al. 2011; Sewell and Fuxman Bass 2018). eY1H assays involve two components: a ‘DNA-bait’ (*e.g.*, a genomic variant, a novel DNA motif, or a repetitive element) and a ‘TF-prey’. DNA-baits are cloned upstream of two reporter genes (*HIS3* and *LacZ*) and integrated into the yeast genome. The DNA-bait yeast strains are then mated with TF-prey strains that express TFs fused to the yeast Gal4 activation domain (AD) to generate diploid yeast containing both bait and prey. If the TF-prey binds to the DNA-bait, the AD will induce reporter expression which can be measured by the conversion of the colorless X-gal to a blue compound (by the β–galactosidase enzyme encoded by *LacZ*), and by the ability of the yeast to grow on media lacking histidine even in the presence of 3-amino-triazole (a competitive inhibitor of the His3p enzyme).

Given that eY1H assays can be used to analyze the DNA-binding activity of more than 1,000 TFs in a single experiment, this framework is particularly well suited to identify the sets of TFs that bind to a DNA region of interest (rather than the sets of DNA regions bound by a TFs as in ChIP-seq). In particular, eY1H assays (and other variations of the assay) have been used to identify the repertoire of TFs that bind to gene promoters and enhancers in humans, mice, nematodes, flies, and plants (Brady et al. 2011; Hens et al. 2011; Gubelmann et al. 2013; Reece-Hoyes et al. 2013; Burdo et al. 2014; Fuxman Bass et al. 2015; Fuxman Bass et al. 2016a). Further, using eY1H assays, we determined altered TF binding to 109 SNVs associated with different genetic diseases including immune disorders, developmental malformations, cancer, and neurological disorders (Fuxman Bass et al. 2015). This pipeline was based on PCR from human genomic DNA using wild type and mutated primers to generate the DNA-baits. As a consequence, the approach used was not suitable to clone and study indels (unless patient DNA samples are used) or novel DNA motifs from footprinting studies. In addition, the eY1H pipeline was not previously adapted to evaluate repetitive DNA elements given that the cloning steps were optimized for unique genomic DNA regions.

Here, we present two novel approaches, one to study repetitive DNA elements, and another to study short DNA sequences (SNVs, indels, and novel DNA motifs) (**Figure 1**). Using the first eY1H approach, we identified 75 TFs that can bind to different Alu sequences present in the human genome. The Alu sequence-binding TFs include nuclear hormone receptor and ETS TFs, some of which are known to bind to these repetitive elements. Using the second approach, we discovered TFs that bind to three novel motifs derived from genome-wide DNase I footprinting by the ENCODE Project that were previously uncharacterized (Neph et al. 2012). Furthermore, we used this method to identify altered TF binding to an 18 bp indel in a T-cell acute lymphocytic leukemia 1 (*TAL1*) super-enhancer and to seven single- or di-nucleotide mutations in the telomerase reverse transcriptase promoter (*TERT*) found in patients with cancer (Horn et al. 2013; Mansour et al. 2014; Huang et al. 2015). Overall, these eY1H-based approaches provide a new toolkit to answer genomic questions that have been challenging to address using current experimental and computational approaches.

**Figure 1.**
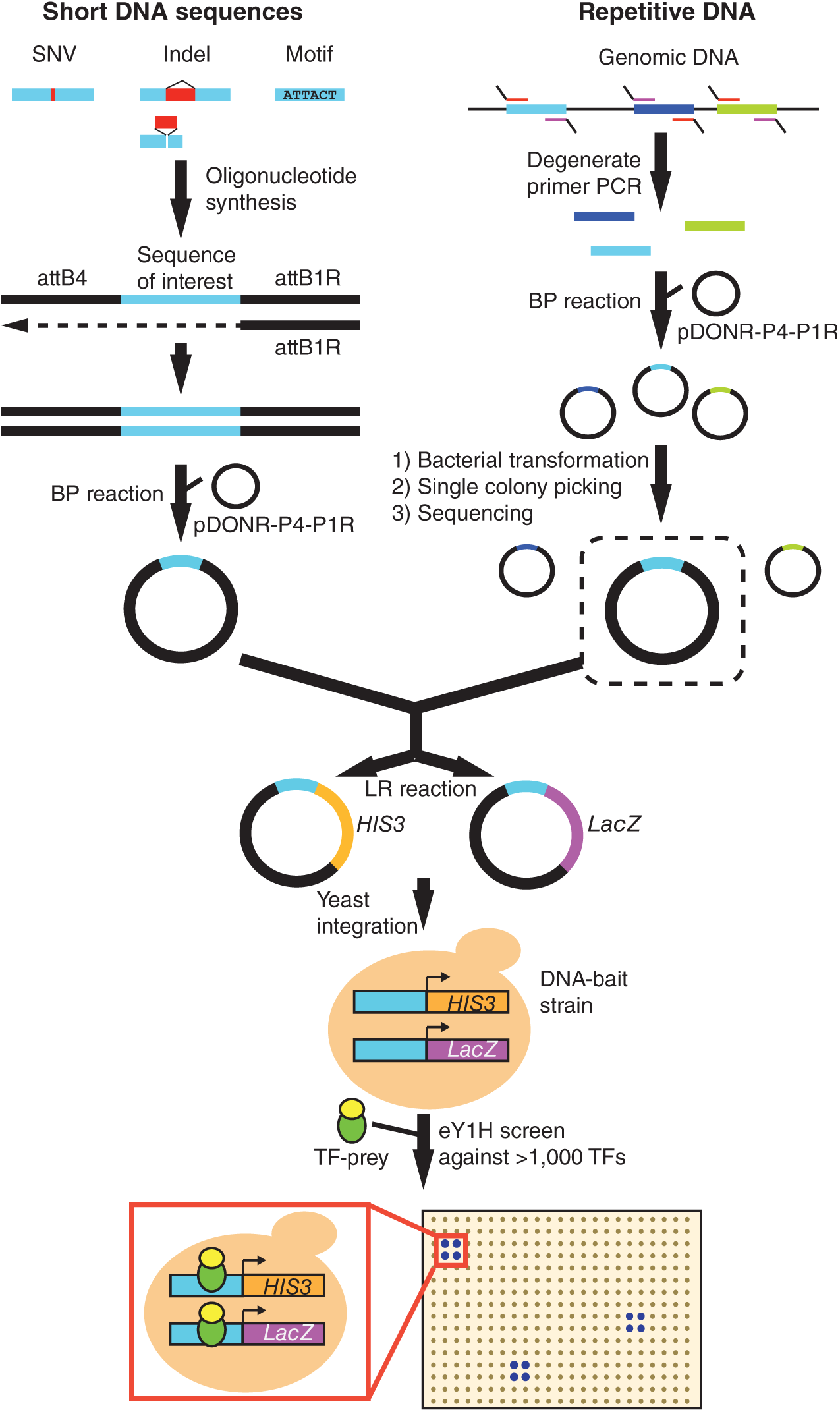
eY1H approaches to test short DNA sequences and repetitive DNA elements. Short DNA sequences are synthesized flanked by the attB4 and attB1R Gateway sites. The reverse strand is synthesized by primer extension (attB1R). The double stranded DNA generated is cloned into the pDONR-P4-P1R vector by Gateway BP reaction. Repetitive DNA sequences are amplified from genomic DNA using degenerate primers flanked by the attB4 (forward) or the attB1R (reverse) sites. The repetitive element DNA library generated is cloned *en masse* into the pDONR-P4P1R vector and individual sequences are selected after bacterial transformation and picking of individual colonies. Both short DNA sequences and repetitive DNA are then transferred into eY1H reporter vectors (*HIS3* and *LacZ*) and integrated into the yeast genome to generate DNA-baits strains. DNA-bait strains are tested for interactions against an array of 1,086 human TF-preys (TFs fused to the yeast Gal4 activation domain) by mating. Interactions are identified by the ability of yeast colonies to grow in the absence of histidine and in the presence of the his3p inhibitor 3-Amino-1,2,4-triazole and to turn blue in the presence of the β-galactosidase substrate X-gal. Interactions are tested in quadruplicate.

## RESULTS

### TF binding to repetitive DNA elements

The binding of TFs to repetitive DNA elements has been challenging to study experimentally given the difficulty of mapping sequencing reads derived from these elements to genomic locations (Rozowsky et al. 2009; Chung et al. 2011). Indeed, most ChIP-seq pipelines currently remove a large fraction of the reads corresponding to repetitive elements (Rozowsky et al. 2009). To illustrate the power of eY1H assays to identify TFs that bind to repetitive elements, we evaluated Alu sequences, a type of short interspaced nuclear element present in more than one million copies in the human genome (Batzer and Deininger 2002). Studying TF binding to these sequences is particularly important given that Alu sequences are embedded within gene promoters, enhancers, and introns, and have been shown to play key roles in gene regulation (Batzer and Deininger 2002; Deininger 2011). In addition, Alu sequences are often silenced by mechanisms that are not fully understood, which could in part be mediated by transcriptional repressors (Humphrey et al. 1996; Kondo and Issa 2003).

To evaluate TF binding to these repetitive elements, we cloned 20 random Alu sequences into our eY1H pipeline (**Figure 1** and **Supplemental Table S1**). This was performed using degenerate primers complementary to the 5’ and 3’ ends of Alu sequences, which allowed us to obtain clones belonging to different Alu families, including the ancestral AluJ, the derived AluS, and the more recent AluY elements (**Supplemental Figure S1**) (Batzer and Deininger 2002). Using eY1H assays, we identified 75 TFs that bind to at least one Alu sequence, and 34 TFs that bind to at least 20% of the 20 Alu sequences tested (**Figure 2A** and **Supplemental Table S2**). Interestingly, Alu sequences are enriched in binding to TFs belonging to the nuclear hormone receptor (NHR), zinc finger DHHC (ZF-DHHC), ETS, and regulatory factor X (RFX) families compared to the array of TFs tested (**Figure 2B**). Of note, we did not detect interactions with NHR or ZF-DHHC TFs for the two AluJ and a subset of AluS sequences tested, suggesting that binding sites for these TFs arose sometime during AluS divergence. Other than this, differences in TF binding between Alu sequences do not seem to cluster by Alu family, likely because of differences in deletions and truncations within family members (**Supplemental Figure S1**).

**Figure 2.**
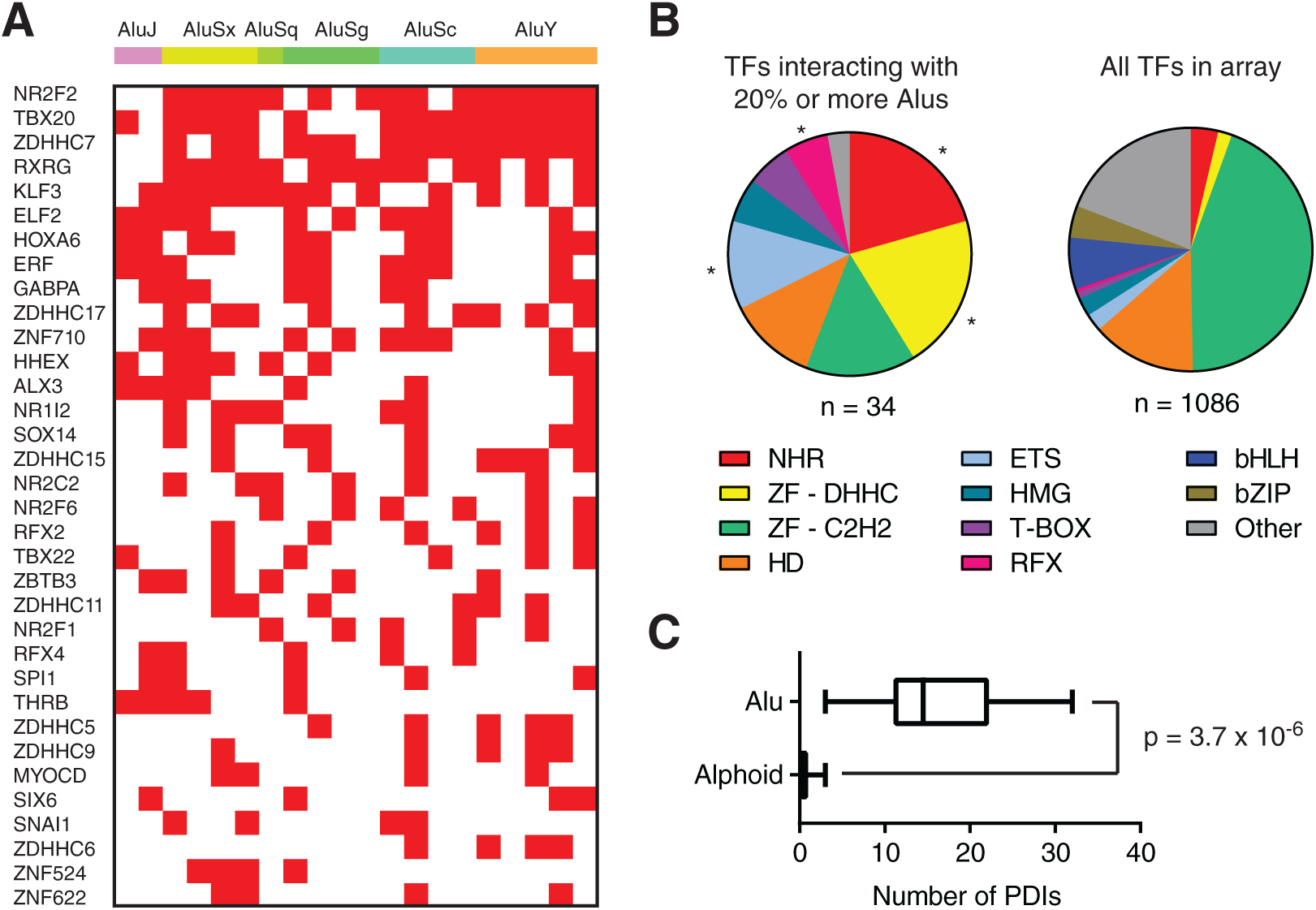
Identification of TFs that interact with Alu sequences. (**A**) TFs that interact with 20% or more of the Alu sequences tested were identified using eY1H assays. TFs are ordered from top to bottom based on the number of Alu sequences they bind. (**B**) Distribution by family for TFs that interact with at least 20% of the Alu sequences tested, compared to the distribution of TFs in the eY1H array. NHR – nuclear hormone receptor, ZF-DHHC – zinc finger DHHC, ZF-C2H2 – zinc finger cys2his2, HD – homeodomain, ETS – E26 transformation specific, HMG – high mobility group, RFX – regulatory factor X, bHLH – basic helix-loop-helix, bZIP – basic leucine zipper domain. *p<0.05 by proportion comparison test after Bonferroni correction. (**C**) Comparison between the number of protein-DNA interactions (PDIs) detected per element for Alu and alphoid sequences. Statistical significance determined by two-tailed Mann-Whitney’s U test.

The widespread TF binding to Alu sequences that we observed is not a general feature of repetitive elements or of eY1H assays, as screening alphoid DNA (*i.e.*, centromeric DNA sequences) only led to marginal TF binding (**Figure 2C**). Indeed, we did not identify any TFs that bound more than 20% of the 12 alphoid sequences tested (not shown), compared to 34 TFs for the Alu sequences. The infrequent binding of TFs to alphoid DNA is consistent with previous findings showing that alphoid DNA recruits multiple centromeric and heterochromatin proteins but not TFs (Buxton et al. 2017). Taken together, our results show that eY1H assays can identify specific TF binding to highly repetitive genomic sequences such as Alu sequences. Whether these TFs globally affect the function of Alu sequences or the function of Alu sequences at specific *loci* remains to be determined.

### Identifying TFs binding to novel DNA motifs

Different experimental methods, including protein-binding microarrays, SELEX, bacterial one-hybrid assays, and ChIP-seq have identified DNA binding motifs for hundreds of human TFs (Noyes et al. 2008; Jolma et al. 2013; Weirauch et al. 2014). However, 289 (out of 683) *de novo* DNA motifs identified by genome-wide footprinting using DNase I by the ENCODE Project (Neph et al. 2012) remain orphan (*i.e.*, no TF has been predicted to bind these motifs). This can stem from the lack of DNA binding motifs for many human TFs (∼50%), from differences between DNA motifs determined *in vitro* and those occupied *in vivo*, from motif quality, from limitations in prediction algorithms, or from DNase I cleavage biases.

To determine whether eY1H assays can identify TFs that bind to some of these orphan DNA motifs, we tested seven well-defined 8 bp DNA motifs identified by DNase I footprints by the ENCODE Project (Neph et al. 2012) that could not be matched to any human TF (**Supplemental Table S3**). To do this, we developed a pipeline using synthesized short oligonucleotides containing the sequence of interest flanked by the Gateway attB4 and attB1R sites for cloning purposes (**Figure 1**). Each DNA motif was tested using three tandem repeats and a two-nucleotide mutated version of the motif as control (**Figure 3A**). For three motifs (UW.Motif.0118, UW.Motif.0146, and UW.Motif.0167) of seven tested, we identified TFs that could interact with the wild type but not with the mutant motifs. For UW.Motif.0118 (GCTGATAA) we found that GATA4, GATA5, and DMBX1 bind to the wild type but not the mutant DNA motif (**Figure 3B**). Indeed UW.Motif.0118 matches the DNA binding motifs for GATA4 and GATA5, which were reported by later publications (**Figure 3C**) (Jolma et al. 2013; Kulakovskiy et al. 2013). More importantly, we found that gene promoters that contain one or more copies of UW.Motif.0118 are enriched in ChIP-seq peaks for GATA4 (**Figure 3I**). DMBX1 can be discarded as a candidate to bind UW.Motif.0118 as its motif matches the junction between two motif copies in the tandem repeat rather than the motif itself (not shown). We also determined that UW.Motif.0146 (ATTTCTGG), but not the mutated motif, binds the zinc finger TF ZBTB26 (**Figure 3D**). The DNA binding motif for ZBTB26 has not yet been determined. Using a position weight matrix prediction algorithm designed for cys2-his2 zinc finger TFs, we predicted a recognition motif based on the amino acid sequence of zinc fingers 1-3, which closely resembles UW.Motif.0146 (**Figure 3E**). Further, we found that gene promoters that contain one or more copies of UW.Motif.0146 are enriched in ChIP-seq peaks for ZBTB26, further validating our approach (**Figure 3I**). Finally, for UW.Motif.0167 (ACAAAAGA) we found that multiple SOX TFs and ZNF646 bind to the wild type but not the mutant motif in eY1H assays (**Figure 3F**). This DNA motif partially matches SOX motifs (**Figure 3G**), while motifs are not available for ZNF646, a protein with 31 zinc fingers according to Uniprot. For several clusters of three ZNF646 zinc fingers, we predicted a CAAAA binding preference, a sequence present in UW.Motif.0167 (**Figure 3H**). This suggests that ZNF646 binds the tandem repeats of the motif used in the eY1H assays rather than a single copy of the motif which are more frequent in the genome.

**Figure 3.**
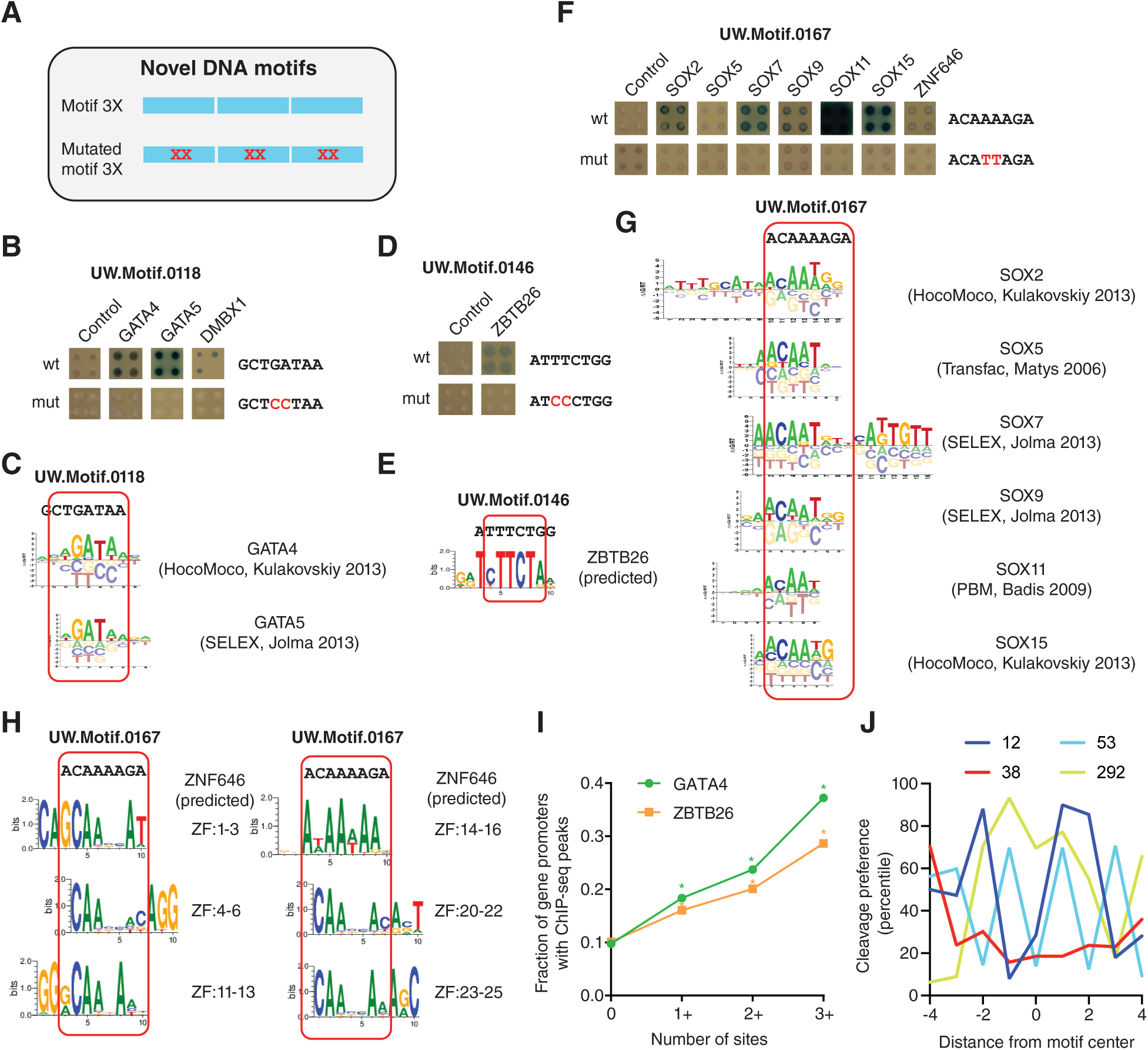
Identification of TFs that bind to novel DNA motifs. (**A**) Motifs were tested by eY1H assays as three tandem copies. Motifs with two mutated bases were tested as controls. (**B, D, F**) eY1H screen of three motifs identified by DNase I footprinting by the ENCODE Project. Tested sequences are indicated. Each interaction was tested in quadruplicate. Control – empty AD-vector. (**C, G**) Alignment of motif logos from CIS-BP to the tested sequences. (**E**) Predicted ZBTB26 motif based on the amino acid sequence of zinc fingers 1-3. (**H**) Predicted ZNF646 motif based on the amino acid sequence of zinc fingers 1-3, 4-6, 11-13, 14-16, 20-22, or 23-25. (**I**) Fraction of genes with ChIP-seq peaks for GATA4 and ZBTB26 in their promoters as a function of the number of UW.Motif.0118 and UW.Motif.0146 motifs, respectively. *p<0.001 vs absence of motif by proportion comparison test. (**J**) Percentile cleavage preference by DNase I at each position for UW.Motif.0012, UW.Motif.0038, UW.Motif.0053, and UW.Motif.0292.

To determine whether the TFs that bind to the orphan DNA motifs are functionally related to their potential respective target genes, we determined the Biological Process Gene Ontology terms associated with genes that contain more than one copy of the motif in their promoter. We found that these genes are associated with similar functions to those determined for the TFs that bind the DNA motifs in eY1H assays (**Supplemental Table S4**). For example, genes that contain more than one copy of the UW.Motif.0118 in their promoters are associated with cardiovascular and neuronal development. This is consistent with the role of GATA4 and GATA5 in heart development and angiogenesis, and the role of GATA4 in neuronal development and function (Lawson and Mellon 1998; Holtzinger and Evans 2007; Walsh and Shiojima 2007; Ang et al. 2016). Similarly, genes that contain more than one copy of the UW.Motif.0167 in their promoters are associated with dendritic spine development, among other biological processes, as are SOX2, SOX5, and SOX11 (Whitney et al. 2014; Hoshiba et al. 2016; Naudet et al. 2018). Altogether, this shows that our approach can identify TFs that bind to uncharacterized motifs, including zinc fingers which are generally difficult to study, and that the TFs identified are functionally related to their potential target genes.

Four motifs (UW.Motif.0012, UW.Motif.0038, UW.Motif.0053, and UW.Motif.0292) out of seven tested (**Supplemental Table S3**) did not produce any interacting TF. This may be due to limitations of eY1H assays including: 1) 25% TFs missing from the human TF array which would not be detected in the screen, 2) TFs that do not fold properly in yeast or fused to AD, and 3) TFs that require post-translational modifications or heterodimerization to bind to DNA (Sewell and Fuxman Bass 2018). Alternatively, some of these motifs may result from sequence bias in DNase I cleavage rather than from nuclease protection by a TF, as has been widely reported (Koohy et al. 2013; Lazarovici et al. 2013; He et al. 2014). Indeed, based on published 6-mer sequence preferences for DNase I (Lazarovici et al. 2013), cleavage within the GAAAAAAA sequence of UW.Motif.0038 is expected to be low (**Figure 3J**). This suggests that the footprints driving UW.Motif.0038 result from a lower ability of DNase I to cleave within this motif rather than from TF protection.

### Identifying altered TF binding to noncoding SNVs and indels

Previously, we used eY1H to identify altered TF binding to noncoding germline variants (Fuxman Bass et al. 2015). That approach used DNA sequences generated by PCR from human genomic DNA as a template and primers containing wild type or mutant sequences to introduce the allele variants (Fuxman Bass et al. 2015). As a consequence, our previous cloning strategy presented several limitations: 1) the requirement of human DNA samples, 2) indels could not be successfully evaluated (unless genomic DNA was obtained from patient samples) as primers containing indels would fail to anneal to the wild type DNA template, and 3) the introduction of unwanted mutations during PCR even when using high fidelity polymerases, thus reducing the efficiency to generate DNA-baits without spurious mutations.

To address these limitations, we leveraged the cloning strategy used for testing novel DNA motifs, and synthesized oligonucleotides containing a short sequence of interest flanked by the attB4 and attB1R gateway cloning sites (**Figure 1**). To establish the optimal oligonucleotide length for testing TF binding to SNVs and indels, we analyzed the motifs present and the published eY1H interactions detected in 168 sequences of 61 bp (Fuxman Bass et al. 2015). We found that the number of motifs detected up to 10 bp from the 5’ and 3’ ends of the sequences is markedly reduced (**Figure 4A**). In addition, we found that the relative fraction of motifs with eY1H interactions varies up to 30% depending on the position, with a minimum at -20 from the attB1R site that increases up to position -4 (**Figure 4A**). Based on these observations, we determined that testing SNVs and indels within ±10 bp of their genomic sequence context (21 bp sequences for SNVs, and 20 bp + indel length for indels) would capture most motifs and provide high sensitivity.

**Figure 4.**
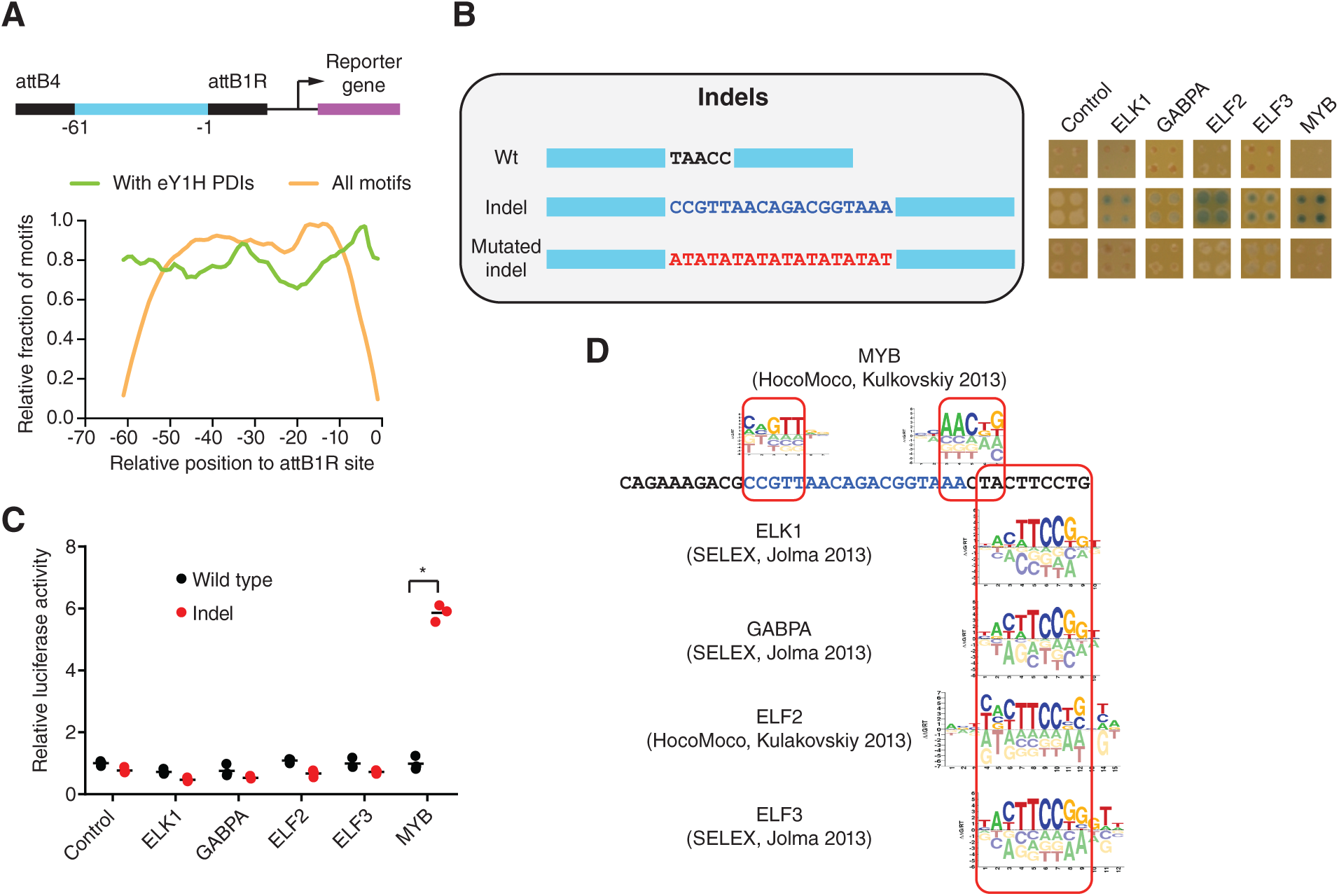
Identification of altered TF binding to a *TAL1* super-enhancer insertion. (**A**) Relative fraction of motifs spanning the indicated position relative to the attB1R site (orange line) and the relative fraction of motifs for which eY1H interactions were detected (green line) for 168 sequences of 61 bp tested by eY1H. Sliding windows of 3 bp were used. (**B**) eY1H screen for an 18bp insertion in a *TAL1* super-enhancer associated with T-cell acute lymphoblastic leukemia. Wild type and an (AT)_9_ sequence replacement (for the 18 bp insertion) were screened as controls. Each interaction was tested in quadruplicate. Control – empty AD-vector. (**C**) Luciferase assays to validate the differential TF interactions with the *TAL1* super-enhancer wild type and insertion alleles. HEK293T cells were co-transfected with reporter plasmids containing the wild type or insertion *TAL1* super-enhancer region cloned upstream of the firefly luciferase reporter gene, and expression vectors for the indicated TFs (fused to the activation domain VP160). After 48 h, cells were harvested and luciferase assays were performed. Relative luciferase activity is plotted as fold change compared to cells co-transfected with the wild type TAL1 super-enhancer construct and the VP160 vector (control). A representative experiment of three is shown. The average of three replicates is indicated by the black line. *p<0.05 by one-tailed log-transformed Student’s t-test with Benjamini-Hochberg correction. (**D**) Motifs obtained from CIS-BP that match the differential TFs identified by eY1H assays.

To evaluate this approach, we tested eight noncoding somatic mutations found in cancer patients: an 18 bp indel in a super-enhancer of *TAL1* and seven single- or di-nucleotide mutations in the *TERT* promoter (**Supplemental Tables S3 and S5**). First, we evaluated whether there is altered TF binding caused by an 18 bp insertion in a *TAL1* super-enhancer found in a patient with T-cell acute lymphoblastic leukemia (Mansour et al. 2014). We detected interactions of TFs ELK1, GABPA, ELF2, ELF3, and MYB with the insertion allele, but not with the wild type sequence or an altered sequence where the 18 bp insertion was replaced by an (AT)_9_ repeat (**Figure 4B**). This control replacing the insertion with a different sequence is important as differences in TF binding between wild type and insertion alleles could arise from differences in distance relative to the minimal promoter of the reporter genes or in nucleosome occlusion. To validate the differential TF-DNA interactions found, we performed reporter assays by co-transfecting HEK293T cells with a reporter plasmid carrying a ∼0.5 kb *TAL1* enhancer sequence containing either the wild type or indel alleles driving luciferase expression and an expression plasmid for the indicated TFs fused to the VP160 (10 copies of VP16) activation domain. The interaction between MYB and the *TAL1* insertion allele was confirmed in human cells by luciferase assays (**Figure 4C**), consistent with a previous study that found that this 18 bp insertion creates a binding site for MYB (Mansour et al. 2014). However, the eY1H interactions between the *TAL1* insertion allele and the ETS factors ELK1, GABPA, ELF2, and ELF3 were not confirmed by the reporter assays (**Figure 4C**), even though these TFs are predicted to bind outside the indel (**Figure 4D**). This difference between eY1H assays (and motif predictions) and luciferase assays may be related to differences in chromatin context (*i.e.*, eY1H assays test interactions within chromatinized DNA *versus* luciferase assays in episomal vectors) or differences in cellular context (*i.e.*, interactions tested in yeast for eY1H assays *versus* human cells in luciferase assays). Additional experiments in the endogenous locus using genome edited cell lines will ultimately determine whether ETS factors affect *TAL1* gene expression caused by the insertion. Regardless of whether the interactions of the altered *TAL1* locus with ETS factors are ultimately validated, it is important to note that eY1H assays narrowed down the follow-up studies to five candidate TFs compared to *in silico* analyses using default settings in the Catalog of Inferred Sequence Binding Preferences (CIS-BP) database (Weirauch et al. 2014) that led to predicting 30 losses and 98 gains of interaction with the *TAL1* super-enhancer insertion (not shown). This is particularly important, given that validation methods such as reporter assays, ChIP, and TF knockdowns are generally low-throughput.

We next applied eY1H assays to test seven single- and di-nucleotide mutations in the *TERT* promoter that lead to telomerase reactivation in different types of cancer including melanoma, bladder cancer, and thyroid cancer (**Figure 5A** and **Supplemental Table S5**). Previously, two *TERT* promoter mutations, -124 G→A and -146 G→A were shown to lead to new interactions with the ETS factor GABPA using reporter assays and TF knockdowns (Bell et al. 2015). Consistent with this, we found that the -146 G→A mutation leads to gain of eY1H interactions with GABPA and ERF, another ETS factor, which were confirmed by motif analysis and luciferase reporter assays, further validating our approach (**Figure 5A**). Interestingly, we found that a two-nucleotide mutation (GG→AA) in the *TERT* promoter at position -138/-139 found in patients with melanoma and bladder cancer (Horn et al. 2013; Wu et al. 2014), for which TF binding was not tested experimentally, leads to gain of interactions with the ETS factors GABPA, ELK1, ELK4, ETV1, ELF2, and ERF (**Figure 5B**). Indeed, the -138/-139 GG→AA mutation creates binding sites for these TFs (**Figure 5C**) but not for other ETS factors such as SPI1 and ETV7 that have slightly different DNA binding specificities (not shown). We found that GABPA, ELK1, ELK4, ETV1, and ERF lead to a stronger activation of the mutant promoter in luciferase reporter assays in human cells, whereas ELF2 did not induce reporter expression (**Figure 5D**). In addition, we found that a -94 A→C mutation in the *TERT* promoter also leads to gain of interactions with GABPA both by eY1H and luciferase assays (**Figure 5A**). Overall, these results suggest that binding sites for GABPA (and other ETS factors) created by single or di-nucleotide mutations at different positions in the *TERT* promoter are responsible for *TERT* reactivation in tumors derived from different cancer patients.

**Figure 5.**
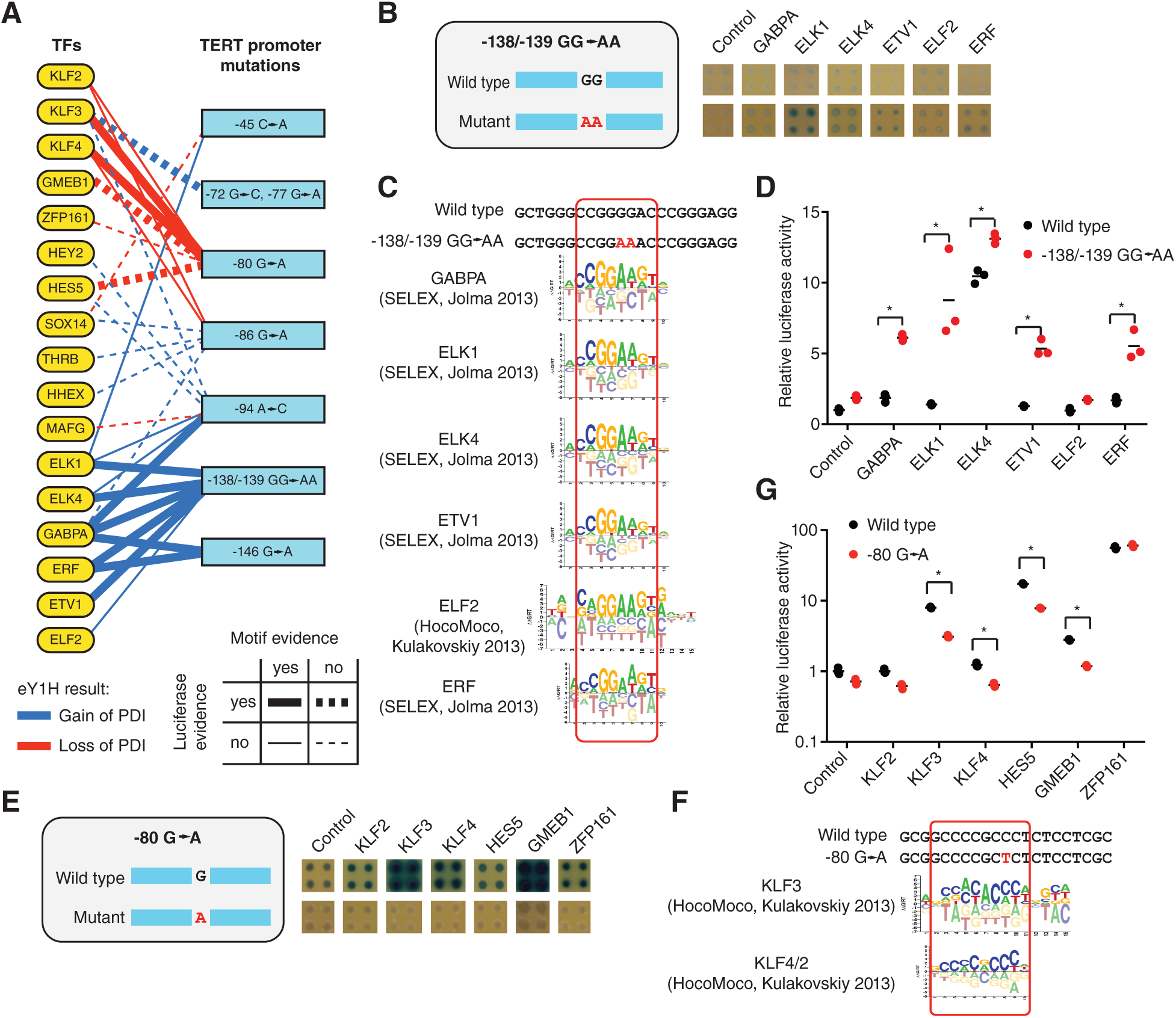
Identification of altered TF binding to single- and di-nucleotide mutations in the *TERT* promoter. (**A**) Gain and loss of protein-DNA interactions (PDIs) with seven *TERT* single- and dinucleodide mutations in the *TERT* promoter were identified by eY1H assays. Mutation position is relative to the initiation codon. Blue line – gain of PDIs, red line – loss of PDIs. Full lines indicate validation by motif analysis. Thick lines indicate validation by luciferase assays. (**B, E**) eY1H screen for wild type and a -138/-139 GG→AA (**B**) or a -80 G→A (**E**) mutation in the *TERT* promoter associated with cancer. Each interaction was tested in quadruplicate. Control – empty AD-vector. (**C, F**) Motifs obtained from CIS-BP that match the differential TFs identified by eY1H assays. (**D, G**) Luciferase assays to validate the differential TF interactions with the TERT promoter alleles. Relative luciferase activity is plotted as fold change compared to cells co-transfected with the wild type TERT promoter construct and the VP160 vector (control). A representative experiment of three is shown. The average of three replicates is indicated by the black line. *p<0.05 by one-tailed log-transformed Student’s t-test with Benjamini-Hochberg correction.

In addition to the gain of interactions with ETS factors, *TERT* promoter mutations can also affect the binding of other TFs. For example, using eY1H assays, we found that a -80 G→A mutation leads to loss of interactions with KLF2/3/4, GMEB1, HES5, and ZFP161 (**Figure 5E**). Loss of interaction with the KLF factors was confirmed by motif analysis and/or luciferase assays, whereas loss of interaction with GMEB1 and HES5 was only confirmed by luciferase assays (**Figure 5A, 5F**, and **5G**). Interestingly, KLF2 and KLF4 have been shown to repress *TERT* expression in normal T cells and lung cancer cells, respectively, although through binding sites that do not overlap with position -80 (Hara et al. 2015; Hu et al. 2016). More importantly, high KLF2/3/4 expression levels are associated with a favorable prognosis in liver, renal, and lung cancer (Hu et al. 2016; Uhlen et al. 2017). Overall, these results suggest that *TERT* reactivation can occur either due to low expression of KLF factors or by mutations in KLF binding sites.

## DISCUSSION

In this study we describe two eY1H approaches to evaluate TF binding to short DNA sequences (*e.g.*, SNVs, indels, and novel DNA motifs) and repetitive elements. Testing these types of sequences has previously been challenging due to limitations in ChIP-seq and DNA motif analyses. For example, motif analyses using CIS-BP (Weirauch et al. 2014) led to the prediction of 128 and 56 differential TF interactions with the *TAL1* super-enhancer 18 bp insertion and the *TERT* -138/-139 GG→AA mutation, respectively. Our approach greatly reduced the number of different TFs that required validation, which is technically important given that mammalian cell reporter assays, ChIP, and TF knockdowns followed by RT-qPCR are generally low-throughput. More importantly, our eY1H approach enables the identification of TF-DNA binding even in the absence of *a priori* TF candidates or a human DNA template as in the case of novel DNA motifs identified by DNase I footprinting assays. For example, we identified TFs that bind to three of seven orphan motifs identified by DNase I footprinting, which is consistent with available or *in silico* predicted motifs, ChIP-seq data, and functional enrichment of target genes. Although some orphan motifs may result from DNase I cleavage bias as previously shown (He et al. 2014) and as we have observed for UW.Motif.0038, our results show that other orphan motifs may indeed result from TF protection.

Our eY1H approach can also be applied to DNA motifs enriched in the regulatory regions of functionally related genes, in particular when these DNA motifs cannot be assigned to any TF. Several factors need to be considered when designing sequences to test motifs by eY1H. First, using multiple tandem copies (three in our case) of a motif increases the sensitivity of the assay. Second, TF interactions may not only occur within the motif but also with the junction between motifs, the junction between the motifs and the Gateway attB4 or attB1R sites, with two consecutive motifs in case of TFs that bind as homodimers or that have more than one domain that recognizes the motif. Thus, it is recommended to also test a mutated version of the motif affecting one or more high information content bases, and to perform motif analysis with the full oligo sequence to exclude TF that do not bind to the motif.

Using eY1H assays we also identified 34 TFs that bind to at least 20% of the repetitive Alu sequences tested. These TFs may be involved in regulating the expression of nearby genes or silencing Alu sequences. Alternatively, Alu sequences may also act as sinks for some TFs, reducing their effective nuclear concentration. However, we did not detect a markedly higher number of sequencing reads matching Alu sequences in ChIP-seq datasets from the ENCODE Project corresponding to Alu-binding TFs than in ChIP-seq datasets corresponding to non Alu-binding TFs (not shown). This could be due to epigenetic silencing of many Alu sequences, which would prevent TF binding to chromatin in human cells in most tissues and conditions. Indeed, most Alu sequences have been found to be enriched in the H3K9me mark and to be actively silenced in somatic tissues (Kondo and Issa 2003; Ward et al. 2013). Nevertheless, thousands of Alu sequences in the human genome contain active histone marks and may be permissive for TF binding which could contribute to the transcriptional control of nearby genes in specific cells and conditions (Deininger 2011; Bouttier et al. 2016). For example, a recent study found that *de novo* ChIP-seq peaks for the H3K4me1 mark in macrophages infected with *Mycobacterium tuberculosis* contain Alu sequences enriched for binding sites of several TFs including ETS and NHR factors, consistent with our findings by eY1H assays (Bouttier et al. 2016). We anticipate that the approach we developed to study TF binding to Alu and alphoid sequences will shed light on the role of other repetitive elements in gene regulation and/or the establishment of heterochromatin.

Overall, the eY1H approaches described here demonstrate their utility for characterizing altered TF binding to different types of genomic variants and for studying the role of TFs in regulating the function of repetitive genomic elements.

## METHODS

### eY1H assays

DNA-baits were generated using different approaches depending on the type of sequence cloned (**Figure 1**). For the repetitive DNA elements, DNA-baits were generated by PCR using human genomic DNA (Clontech) as a template, Platinum Hifi Taq polymerase (ThermoFisher), and degenerate primers complementary to different family members of Alu elements (Alu-Fw and Alu-Rv, **Supplemental Table S3**) or different variations of alphoid DNA (Alphoid-Fw and Alphoid-Rv, **Supplemental Table S3**) (**Figure 1**). These primers include the attB4 and attB1R sequences for Gateway cloning. The PCR cycle involved an initial denaturation step of 2 min at 94°C, 35 cycles of 30 sec at 94°C, 15 sec at 58°C, and 75 sec at 72°C, followed by a final extension for 7 min at 72°C. The random libraries containing the Alu sequences or the alphoid DNA were cloned into the pDONR-P4-P1R vector (ThermoFisher) by Gateway cloning using the BP Clonase II (ThermoFisher) and transformed into DH5α bacteria. Individual colonies were picked and sequenced to identify the sequences cloned (**Supplemental Table S1**). Each sequence was then transferred to the pMW#2 and pMW#3 vectors (Addgene) using the LR Clonase II (ThermoFisher), upstream of two reporter genes (HIS3 and LacZ). Both reporter constructs were integrated into the Y1HaS2 yeast strain (Reece-Hoyes et al. 2011) genome by site-specific recombination to generate chromatinized DNA-bait strains as previously described (Fuxman Bass et al. 2016b; Fuxman Bass et al. 2016c). The DNA-bait yeast strains were then sequenced to verify the identity of the yeast integrants.

DNA-baits corresponding to genomic variants (SNVs and indels) and novel DNA motifs were synthesized as oligonucleotides (ThermoFisher) flanked by the attB4 and attB1R sequences for cloning using the Gateway recombination system (**Figure 1** and **Supplemental Table S3**). Double-stranded oligonucleotides were generated by primer extension using Taq polymerase (ThermoFisher) and a primer complementary to the attB1R (**Supplemental Table S3**) site using an initial denaturation step of 3 min at 95°C, ten cycles of 30 seconds at 55°C and 30 seconds at 72°C, followed by one cycle of 5 min at 72°C. The double-stranded oligonucleotides were then cloned into the pDONR-P4-P1R (ThermoFisher) by Gateway cloning, transferred to the pMW#2 and pMW#3 vectors, and then integrated into the Y1HaS2 strain genome.

DNA-bait strains were mated with an array of yeast strains expressing 1,086 human TF-preys using a Singer RoToR robotic platform, as previously described (Reece-Hoyes et al. 2011; Fuxman Bass et al. 2015). This TF array includes members from all major human TF families (Fuxman Bass et al. 2015). Each interaction was tested in quadruplicate, and only interactions detected with at least two colonies were considered positive (**Figure 1**). As previously observed, ∼90% of interactions were detected by all four replicates (Reece-Hoyes et al. 2011; Fuxman Bass et al. 2015; Fuxman Bass et al. 2016a).

### Transient transfections and luciferase assays

TF interactions with noncoding alleles were validated by luciferase assays in HEK293T cells. Given that testing the noncoding alleles in the short sequence context (20-40 bp) used in eY1H assays led to luciferase activity barely above background levels (not shown), DNA-bait luciferase reporter clones were generated corresponding to the ∼0.5 kb genomic sequence surrounding the noncoding alleles (**Supplemental Table S6**). Wild-type sequences were generated by PCR using human genomic DNA (Clontech) as a template, Platinum Hifi or SuperFi Taq polymerases (ThermoFisher), and primers surrounding the noncoding alleles. Mutant sequences were generated from the wild-type sequence by PCR stitching using primers that contain the mutated nucleotide. DNA-bait luciferase reporter clones were generated by cloning the noncoding regions upstream of the firefly luciferase into a Gateway compatible vector generated from pGL4.23[luc2/minP] (Fuxman Bass et al. 2015). TF-prey clones were generated by Gateway cloning the TF coding sequence into the pEZY3-VP160 vector where TFs are fused to ten copies of the strong transcriptional activator VP16 (Carrasco Pro et al. 2018).

HEK293T cells were plated in 96-well opaque plates (∼1 x 10^4^ cells/well) 24 hours prior to transfection in 100 µl DMEM supplemented with 10% FBS and 1% Antibiotic-Antimycotic 100X (ThermoFisher). Cells were transfected with Lipofectamine 3000 (ThermoFisher) according to the manufacturer’s protocol using 20 ng of the DNA-bait pGL4.23[luc2/minP] luciferase reporter vector, 80 ng of the TF-pEZY3-VP160 vector, and 10 ng of renilla luciferase control vector pGL4.74[hRluc/TK] (Promega). The empty pEZY3-VP160 vector co-transfected with the recombinant firefly luciferase plasmid was used as a normalization control. Fourty-eight hours after transfection, firefly and renilla luciferase activities were measured using the Dual-Glo Luciferase Assay System (Promega) according to the manufacturer’s protocol. Non-transfected cells were used to subtract background luciferase activities, followed by normalizing firefly luciferase activity to renilla luciferase activity.

### ChIP-seq analysis

ChIP-seq peaks for GATA4 (ENCSR590CNM) and ZBTB26 (ENCSR229DYF) were obtained from the ENCODE Project (Consortium 2012). A peak was assigned to a gene promoter if the midpoint of the peak was located within -2 kb to +250 bp of the transcription start site (according to Gencode v19). In addition, promoter positions overlapping with coding regions were excluded from this analysis. The midpoint of peaks overlapping with promoter regions was calculated using the intersect option from BEDTools (Quinlan and Hall 2010). The fraction of genes with GATA4 and ZBTB26 peaks in their promoters was calculated as a function of the number of UW.Motif.0118 and UW.Motif.0146 motifs, respectively.

### Prediction of DNA binding motif for zinc finger proteins

DNA motif prediction for the Cys2-His2 zinc finger TFs ZBTB26 and ZNF646 was performed using an expanded support vector machine model available at http://zf.princeton.edu/fingerSelect.php (Persikov and Singh 2014). For ZBTB26, zinc fingers 1-3 were used for predictions. For ZNF646, zinc fingers 1-3, 4-6, 11-13, 14-16, 20-22, and 23-25 were used for predictions.

### Gene ontology enrichment

For each of the DNA motifs derived from DNase I footprints, the Biological Process Gene Ontology enrichment was determined using the Gene Ontology Consortium enrichment analysis tool using all human genes as background (http://geneontology.org/page/go-enrichment-analysis) for genes with at least two motifs in their promoter regions. For each cluster of terms, only the term with the highest fold enrichment (>2) was considered. For each of the TFs found to bind to the DNA motifs in eY1H assays (except ZBTB26 and ZNF646 for which no or few publications are available), the association between the TF and the gene ontology terms was annotated (**Supplemental Table S4**).

### DNase cleavage preference

DNase I cleavage expected preferences within each 8 bp motif was calculated using 6-mer cleavage frequencies determined in ref(Lazarovici et al. 2013). For cleavage sites at positions -4, -3, and -2 from the center of the motif, the average cleavage frequency was calculated between 6-mers NNNXXX, NNXXXX, and NXXXXX, respectively, where N is any nucleotide and X correspond to a nucleotide in the motif. For cleavage sites at positions 2, 3, 4 from the center of the motif, the average cleavage frequency was calculated between 6-mers XXXXXN, XXXXNN, and XXXNNN, respectively. The percentile cleavage preference at each position was determined by comparing to all the averaged frequencies at their respective positions.

### Selection of oligonucleotide length to test SNVs and indels

To determine the optimal oligonucleotide length to evaluate TF binding to SNVs and indels, the motifs present and the published eY1H interactions detected in 168 sequences of 61 bp were analyzed (Fuxman Bass et al. 2015). For each position relative to the attB1R cloning site (which is proximal to the minimal promoter of the reporter genes), the number of motifs spanning the indicated position and the fraction of motifs with the corresponding eY1H interaction was calculated. Reported values are relative to the maximum and correspond to sliding windows of 3 bp (except at positions -1 and -61).

## DATA ACCESS

The protein-DNA interactions from this publication have been submitted to the IMEx (http://www.imexconsortium.org) consortium through IntAct (Orchard et al. 2014) and assigned the identifier IM-26689.

### ACKNOWLEDGMENTS

We thank Drs. Trevor Siggers and Thomas Gilmore for critically reading the manuscript. This work was supported by the National Institutes of Health Grants R00-GM114296 and R35-GM128625 to J.I.F.B.. J.A.S. was supported by NIH training grant 5T32HL007501-34 and M.M by a National Science Foundation REU [BIO-1659605].

## AUTHORS CONTRIBUTIONS

J.I.F.B., J.A.S., S.S., E.F., and M.M. performed eY1H assays. S.S. and C.S.S. performed luciferase assays. J.I.F.B. and S.C.P. performed the data analyses. J.I.F.B. conceived the project and wrote the manuscript. All authors approved the content of the manuscript.

## DISCLOSURE DECLARATION

The authors declare no competing interests.

