## Supplemental Figure S1 for "Discovering human transcription factor interactions with genetic variants, novel DNA motifs, and repetitive elements using enhanced yeast one-hybrid assays"

### Supplementary Figure S1

```
Alu_101-2       CGCCCGTAATCCTAGCACTTTCCAAGGCCGAGGTGGTGGGAGGATTGCTTGAACCCAGGA 60
Alu_92-1        TGCCTATGATCCCAGCACTTTGGGAGGCCAAGG---CAGGCAGATTGCTTGAACCCAGAT 57
Alu_87-1        TGCGTGTAATCCCAGCACTTTGGGAGGCCGAGA---AGGGAGAA--TCACAAGGTCAGGA 55
Alu_6-2         TGCCTGTAATCCAAGCACTTTCGGAGGCCAAGG---CAGGTGGA--TCACGAGGTCAGGA 55
Alu_112-3       CACCTGTAATCCCAGCACCTTGGCAGGCTGAGG---CAGGCAGGTCACCTGAATTCAGGA 57
Alu_100         TGCTTGTAATACCAGGACTTTGGGAGGCCAAGT---GGGGCAGATCGCTTGAGGTCAGGA 57
Alu_207-1       CGCCTGTAATCCCAGCACTTTGGGAGGCCAAGG---TGGGCGGA--TCACAAGGTTGGGG 55
Alu_110-1       CGTCTGTAATCTCAGCACTTTGGGATGCCAAGG---TGGGTAGATCACCTGAGGTCAGGA 57
Alu_111-2       TGCCTGTAATCCCAGCCTTTTGGGAGGCCGAAG---AGAGCGGA--TCACAAAGTCAAGA 55
Alu_33-2        CAACTGTAATCCCAGCAACGTGGGAAGCCGAGG---TGGGCAGA--TCACAAGGTCAGGA 55
Alu_205-1       CACCTGTAATCCCAGCACTTTGGGAGGCTGAGG---CGGGCGGA--TCACAAATTCAAGA 55
Alu_204-1       CGCCTGTAATTCCAGCACTTTGGGAGGCTGAGG---CAGGTGCA--TCACAAGGTCAAGA 55
Alu_133-1       CGCCTGTAATCCCAGCACTTTGGGAGGCCGAGG---CGGGCAGA--TCACGAGGTCAGGA 55
Alu_4-1         TGCCTGTAATCCCAGCACTTTGGGAG-GCCAAG---TGGGCAGA--TCATGAGGTCAGGA 54
Alu_105-1       CACCTGTAATCCCAGCACTTTAGGAGGCCAAGG---AGGGCGGA--TCACGAGGTCAGGA 55
Alu_22-1        TGCCTGTAATCCTAGCACTTTGGGAGACCAAGG---AGGGCAGG--TCATGAGGTCAGGA 55
Alu_72-1        CGCCTGTAATCCCAGCACTTTGGGAGGCCGAGG---CGGGCGGA--TCACGAGGTCAGGA 55
Alu_78-2        CACCTGTAATCCCAGCACTTTGGGAGGCTGAGG---AGGGCAGA--TCACAAGGTCAGGA 55
Alu_142-2       TGCCTGTAATCCCAGAACTTTGGGAGGCTGAGG---TGGGCAGA--TCACGAGGTCAGGA 55
Alu_74-1        TGCTTGTAATCCCAGCACTTTGGGAGGTCAAGG---CAGGCAGATCACCTGAGGTCAGGA 57
                      * **   **     *   *     *        *       *   *

Alu_101-2       GTTCAAGATCATCTTGGAAAACAAAGTGAGACTCCATCTTTACAAAA------------- 107
Alu_92-1        GTTTGAGACCAGCCTGGGCAACATGGCAAAACCCCATCTTTACAAAATGTACAGAAAATA 117
Alu_87-1        GTTCAAGACAAGCCTGACCAATATGTTGAAACCCTATCTCCACTAAAAAAAATGCAAACA 115
Alu_6-2         GTTCGAGACCAGCCTGGCCAAGATGGTGAAACTCAACCCCGTCTCTACTAAAATAC---- 111
Alu_112-3       GTTAGGGACCAGCCTGGCCAACATGGCAAAACCCTATCTCTACTA--AAAATACAA---- 111
Alu_100         GTTCGAGACCAGCCTGGTCAATATGGTGAAATCCCGTCTCTACTA--AAAATAGAA---- 111
Alu_207-1       GTTCGAGACCAGCCTGGCTAACATGGTGAAACCCTGTCTCTACAA--AAAATACAA---- 109
Alu_110-1       GTTCGAGACCAGCCTGCCCAACATGGTGAAACCCCGTCTCTACTA--AAAATACAA---- 111
Alu_111-2       GATAGAGACCATCCTGGCTAACACGGTGAAACCCCATCTCTACTA--AAAGTACAA---- 109
Alu_33-2        GATCGAGACCATCCTGGCCAACAAGGTGAAACCCCGTCTCCACTA--AAAATACAA---- 109
Alu_205-1       GATCAAGACCATCCTGGCCAACATGATAAAACCCCGTATCTAGTA--AAAATACGA---- 109
Alu_204-1       GATCGAGGCCATCCTGGCCAACATGGTGAAACTCCGTCTCTACTA--AAAATACAA---- 109
Alu_133-1       GATCGAGACCATCCTGACCAACATGGTGAAACCCCGTCTCTACTA--AAAATACAA---- 109
Alu_4-1         GATCGAGACTATCCTGGCTAACATGGTGAAACCCCATCTCTACTA--AAAATACAA---- 108
Alu_105-1       GATCAAGACCATCCTGGCTAACACGGTGAAACCCTGTCTCTACTA--AAAATACAA---- 109
Alu_22-1        GATCAAGACCATCCTGGCTAACACGGTGAAACCCCGTCTCTACTA--AAAATACAA---- 109
Alu_72-1        GATCGAGACCATCCCAGCTAAAACGGTGAAACCCCGTCTCTACTA--AAAATACAA---- 109
Alu_78-2        GTTCAAGACCAGCTTGGCCAACATGGTGAAACCCCGTCCCTACTA--AAAATACAA---- 109
Alu_142-2       GTTTGAGACTAGCCTGGCCAATATGGTGAAACCCCGTCTCTACTA--AAAATACAA---- 109
Alu_74-1        GTTCGAGACCAGCCTGGCCAACATGGTGAAACCCCATCTCTACTA--AAAATACAA---- 111
                * *   *   * *      ** *     * *  *

Alu_101-2       ------ATTTTAAAAAGTTAGCTTGCGATGGTGGTGTGTACATGTGTTCTCAGCTACTTG 161
Alu_92-1        AATAAAAATAAAAATAAATAGCTAGGCATGGTGGTGCGTGCCTGTGGTCCCAGCTACTCC 177
Alu_87-1        A---AATACAAAAAAAATTAGCTGGGCGTGGTGGCGCATGCCTGTAATCCCAGCTACTCA 172
Alu_6-2         ------------AAAAATTAGCCAGGCGCGGTGGCGGGCAACTGTAATCCCAGTTACTCG 159
Alu_112-3       ------------A--AATTAGTTGGGTGTGGTAGCTGGGACCTCTAATCCCAGCTACTCG 157
Alu_100         ------------A--AATTAGCTGGGCGTGGTGGTGGGCACCTGTAATCCCAGCTACTTG 157
Alu_207-1       ------------AA-ATTAGCCAGGCTGTAGTGGCGCGTGCCTGTAATCCCAGCTACTCG 156
Alu_110-1       ------------A--AATTAGCCAGCTGCGGTGGTGTGCACCTGTTGTCCCATCTACTT- 156
Alu_111-2       ------------AA--AATTGGCCGGGCGTGGTAGCGTCACCTGTAGGCCCAGCTACTCG 155
Alu_33-2        ------------A--AATTAGCTGAGCATGGCGGCGCACACCTGTAGTCCCAGCTACTCG 155
Alu_205-1       ------------AA-AATTAGCTGGGTGTGGTGGTGTGCACCTGTAGTCCCAGGTACTCG 156
Alu_204-1       ------------A--AATTAGCTAGGCATGGTGGTGTGCACCTGTAGTCCCAGCTACTCA 155
Alu_133-1       ------------AAAAATTAGCCAGGCGTGGTGGTGGGCGCCTGTAGTTCCAGCTGCTCA 157
Alu_4-1         ------------AA-AATTAGCCGGGCATGGTAGCGGGCGCCTATAGTCTCAGCTACTCG 155
Alu_105-1       ------------AA-AATTAGCCAGGCATGGTGGCAGGCACCTGTAGTCCCAGCTACTTG 156
Alu_22-1        ------------AA-AATTAGCTGGGCGTGGTGGCAGGCGCCTGTAATCCCAGCTACTCT 156
Alu_72-1        ------------AA-AATTAGCCGGGCGTAGTGGCGGGCGCCTGTAGTCCCAGCTACTTG 156
Alu_78-2        ------------A--AATTAGCCGGGCGTGGTGGTGCATGCCTGTAATCCCAGCTTCTTG 155
Alu_142-2       ------------A--AGTTAGCCAGGTGTGATGGCATGCGCTTGTAGTCCCAGCTACTCG 155
Alu_74-1        ------------A--AATTAGCAGGGCGTTGTGACATGTGCCTGTAATCCCAGGTACTTG 157
                            *                             * *     **  * **

Alu_101-2       GGAGGCTCAGGTGTGAGGATCACTTCAGCCCAGAAGGTTGAGGCTGCAGTGAGCCAGGGA 221
Alu_92-1        AGAGACTGAGGTGGGAGGATCACCTGAGCCCAGGAGGTAGAGGTTGCAGTGAGCCACTGC 237
Alu_87-1        GGAGGCTGAGGCAGGTGAGCCGAGATTGTGCCA--------------------------- 205
Alu_6-2         GGAGGCTGAGGCAGGAGAATCGCTTGAACCTGGGAGGTGGAGGTTGCAGTGAGCCAAGAT 219
Alu_112-3       GGAGGCTGAGGTGGGAGAATCACTTGAACCCGGGAGGCAGAGGTTGCAGTGAGCCGAAAT 217
Alu_100         GGAGGCTGAGGCAGGAGCATCGCTTGAACCCAGGAGTTGGAGGTTGCAGTGAAATCCTGC 217
Alu_207-1       GGAGGCTGAGGCAGGAGAATTGCTTGAACTAGGGAGGCAGAGGCTGCAGTGAGCCAAGAT 216
Alu_110-1       GGGGGCTGAGGCAGGAGAATCGCTTGAACCTGGGAGGCGGAGGTTGCAGTGAGCTGAGAT 216
Alu_111-2       GGAGGCTGAGGCAGGAGAATCACTTGAACCCAGGAGGTGGAGGTTGCAGTGAGCTGAGAT 215
Alu_33-2        GGAGGCTGAGGCAGGAGAATCGCTTGAGCCCGGGAGGTGGAGGTTGCAGTGAGCCAAGAT 215
Alu_205-1       GGAGGCTGAGGCAGGAGAATCGCTTGAACTTGGGAGGCAGAGGTTGCGGTGAGCCGAGAT 216
Alu_204-1       GGAGGCTGAGGCAAGAGAATTGCTTGAACCCAGGAGGCGGAGGTTGCAGTGAGCCGAGAT 215
Alu_133-1       GGAGGCTGAGGCAGGAGAATGGTGTGAACCCGGGAGTCGGAGTTGGCAGTGAGCCGAGAT 217
Alu_4-1         GGAGGCTGAGGCAGAAGAATGGCGTGAACCTGGGAGGCGAAGCTTGCAGTGAGCCAAGAT 215
Alu_105-1       GGAGGCTGAGACAGGAGAATGGTGTGAACCGGG-AGGCGGAGCTTGCAGTGAGCCGAGAT 215
Alu_22-1        GGAGGCTGAGGCAGGAGAATGGCGTGAACCTGGGAGGCGGAGCTTGCAGCGAGCCAAGAT 216
Alu_72-1        GGAGGCTGAGGCAGGAGAATGGCGTGAACCCGGGAGGCGGAGCTTGCAGTGAGCCGAGAT 216
Alu_78-2        GGAGCCTGAGGCAGGAGAATCATTGGAACCCCGGAGGTGGAGGTTGCAGTGAGCCAAGAT 215
Alu_142-2       GGAGGCTGAGGCAGGAGAATCACTTGAACCCGGGAGGCGGAGGTTGCAGTGAACTGAGAT 215
Alu_74-1        GGAGTCTGAGGCAGGAGATTCACTGGAACCTGGGAGGTAGAGGTTGCAGTGAGTTGAGAT 217
                 * * ** **      *

Alu_101-2       TGGCGCCACTGCTCTCCAGCC----------------------       242
Alu_92-1        ACTCCAGCC----------------------------------       246
Alu_87-1        -------CTGC-ACTCCAGCC----------------------       218
Alu_6-2         CGCTGCACTCT-AGCC---------------------------       234
Alu_112-3       CACACACACCA-TTGCACTCCAGCC------------------       241
Alu_100         CATTGCACTCC-AGCC---------------------------       232
Alu_207-1       CGCACCACTGT-GCTCAAGATTGTACCAGTGCACTCCAGCC---       256
Alu_110-1       CATGCCACTGC-ACTCCAGCC----------------------       236
Alu_111-2       TGCGCCAAAGC-ACTCCAGCC----------------------       235
Alu_33-2        TGTGCCACTGA-ACTCCAGCC----------------------       235
Alu_205-1       TGCACCACTGC-ACTCCAGCC----------------------       236
Alu_204-1       CTCACCACTGC-ACTCCAGCC----------------------       235
Alu_133-1       CGCGCCACTGC-ACTCTAGCC----------------------       237
Alu_4-1         CGTGCCACTGC-ACTCCAGCC----------------------       235
Alu_105-1       TGCGCCAGTGC-ACTCCAGCC----------------------       235
Alu_22-1        CGCACCACTGC-ACTCCAGCC----------------------       236
Alu_72-1        CCCGCCACTGC-ACTCCAGCC----------------------       236
Alu_78-2        CATACCACTGT-ACTCCAGCCTGGGCGACAGGCAAGACTCCATC       258
Alu_142-2       TGCACCACTGC-ACTCCAGCC----------------------       235
Alu_74-1        TGCACCACTGC-ATTCCAGCC----------------------       237
```

AluJ | AluSq | AluSc
AluSx | AluSg | AluY

**Supplemental Figure S1: Alu element sequence alignment.** Alu sequences were aligned using Clustal Omega. Colors indicate the Alu element family.
